# Central and peripheral clocks synergistically enhance circadian robustness, flexibility and performance of cardiac pacemaking

**DOI:** 10.1101/2023.08.07.552251

**Authors:** Pan Li, Jae Kyoung Kim

## Abstract

The strong circadian (∼24h) rhythms in heart rate (HR) are critical for flexible regulation of cardiac pacemaking function throughout the day. While this circadian flexibility in HR is robustly sustained in diverse conditions, it declines as the heart ages, accompanied by reduced maximal HR performance. The intricate regulation of circadian HR patterns involves the orchestration of sympathetic and parasympathetic nervous activities (SNA and PNA) alongside local circadian rhythmicity (LCR) within the heart. However, their intricate interactions that sustain the resilience and adaptability of circadian rhythms, as well as the mechanisms that underpin their deterioration during the aging process, remain enigmatic. To address these questions, we developed a mathematical model describing autonomic control and LCR in sinoatrial nodal cells (SANC) that accurately captures distinct circadian patterns in adult and aged mice. Our model underscores the indispensable synergy among SNA, PNA, and LCR in preserving circadian flexibility, robustness, and performance in SANC. SNA predominantly enhances SANC robustness and performance, while PNA primarily drives SANC flexibility, complemented by LCR and SNA. LCR acts as a booster, further enhancing SANC flexibility and performance. However, the delicate balance of this synergy is disrupted with age, resulting in diminished SANC performance and flexibility. Specifically, age-related impairment of PNA selectively dampens SANC flexibility while ion channel remodeling disrupts all SANC functions. Our work shed light on their critical synergistic interactions in regulating time-of-day cardiac pacemaking function and dysfunction, which may help to identify potential therapeutic targets within the circadian clock for the prevention and treatment of cardiac arrhythmias.

**Author Summary:** The mammalian heart relies on the sinoatrial node, known as the cardiac pacemaker, to orchestrate heartbeats. These heartbeats slow down during sleep and accelerate upon waking, in anticipation of daily environmental changes. The heart’s ability to rhythmically adapt to these 24-hour changes, known as circadian rhythms, is crucial for flexible cardiac performance throughout the day, accommodating various physiological states. However, with aging, the heart’s circadian flexibility gradually weakens, accompanied by a decline in maximal heart rate. Previous studies have implicated the involvement of a master circadian clock and a local circadian clock within the heart, but their time-of-day interactions and altered dynamics during aging remain unclear. In this study, we developed a mathematical model to simulate the regulation of sinoatrial nodal cell pacemaking function by the master and local circadian clocks in adult and aged mice. Our results revealed distinct roles played by these clocks in determining circadian patterns of sinoatrial nodal cells and shed light on their critical synergistic interactions in regulating time-of-day cardiac pacemaking function and dysfunction.

## Introduction

The mammalian heart exhibits robust circadian rhythms in various cardiac function indices, such as heart rate (HR) and electrocardiogram (ECG) waveforms (1). During sleep, HR slows down, accompanied by prolongation of QRS duration and QT interval, while it accelerates upon waking, indicating circadian variations in the electrical properties of the sinoatrial node and ventricular tissue (2). In addition, cardiac arrhythmic phenotypes also show distinct circadian patterns. For instance, while bradyarrhythmias and Brugada syndrome are more prevalent at night, ventricular fibrillation and sudden cardiac death are more common in the morning (2, 3).

These circadian rhythms in cardiac physiology and pathology are regulated by the master circadian clock located in the suprachiasmatic nucleus (SCN)(3). The master clock influences the firing frequency of the sinoatrial node through the autonomic nervous system (ANS), thereby modulating HR through sympathetic and parasympathetic stimulation (Fig 1) (4). Specifically, sympathetic nervous activities (SNA) increase HR by activating G-protein-coupled receptors (GPCR), which in turn activate adenylyl cyclase (AC). Then, AC activation catalyzes the formation of the cyclic adenosine monophosphate (cAMP), which is a ubiquitous second messenger that activates protein kinase A (PKA). This AC-cAMP-PKA signaling cascade leads to the phosphorylation of various targets, such as the L-type Ca^2+^ channel (I_CaL_), Ryanodine receptor, and sarco/endoplasmic reticulum (SR) Ca^2+^ ATPase (SERCA) (Fig 1; yellow dot) (5). In contrast, parasympathetic nervous activities (PNA) decrease HR by activating the muscarinic K^+^ current (I_KACh_) and inhibiting the AC-cAMP-PKA signaling pathway (Fig 1; blue dot) (6). A recent experimental study in immobilized anesthetized mice suggests that PNA may have a dominant role in circadian HR variations compared to SNA (7).

**Fig 1.**
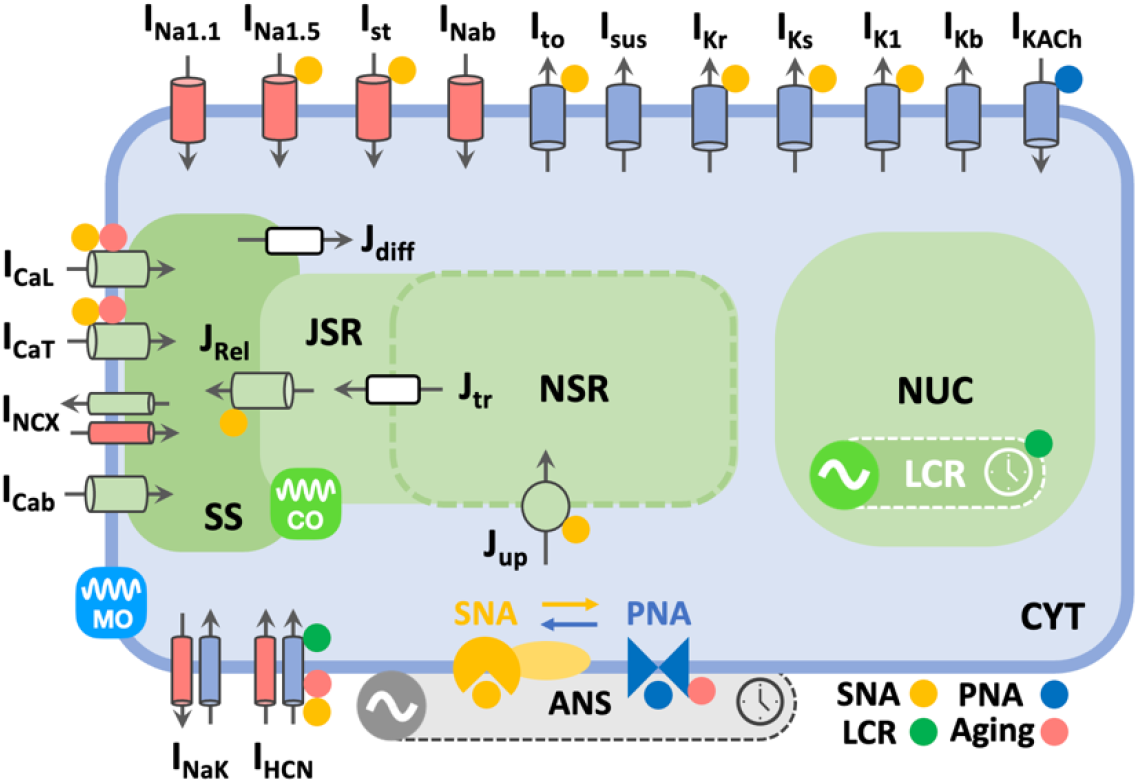
Model schematic for autonomic control and LCR in mouse SANC. During each cardiac cycle (∼ a hundred milliseconds), the membrane oscillation (MO) is primarily driven by ion channels that control the flow of ions across the cell membrane, e.g., I_HCN_ that promotes membrane depolarization, inward I_CaT_, I_CaL_, and Na^+^ currents that trigger spontaneous AP. On the other hand, the Ca^2+^ oscillation (CO) involves Ca^2+^ influx via I_CaT_ and I_CaL_ that triggers intracellular Ca^2+^ release from the SR Ca^2+^ store into the cytosol, and Ca^2+^ removal through SERCA (Jup) and the Na^+^-Ca^2+^ exchanger current (I_NCX_). Over the course of a day and night cycle (∼24hr), SANC pacemaking activities are tightly regulated by both the ANS and LCR to generate circadian variations in firing rates (FR). The ANS is regulated by the master circadian clock - SCN. Simultaneous SNA (orange dots) and PNA (blue dots) co-modulate a diverse range of subcellular targets in SANC and exhibit non-additive effects via AC-cAMP-PKA dependent or independent pathways (6). LCR (green dots) within the SANC nucleus (NUC) leads to circadian variations in the expression levels of ion channels, e.g., I_HCN_. In aged mice, SANC pacemaking function is weakened by aging-dependent ion channel remodeling of I_CaT_, I_CaL,_ and I_HCN_, and PNA impairment (pink dots) (14, 15).

Besides the ANS, circadian rhythms in HR are also influenced by LCR (Fig 1; green dot), exemplified by diurnal variations in the expression level of the hyperpolarization-activated cyclic nucleotide–gated (HCN) channel (I_HCN_; or I_f_) (8). Prior experimental studies showed that circadian HR rhythms are lost in SCN-lesioned mice, while they are preserved in mice with local circadian disruption, suggesting a primary role of the ANS and a secondary role of LCR in regulating HR circadian rhythms (9-11). However, under ANS blockade conditions, further studies demonstrate the significant contribution of LCR in promoting diurnal variations in intrinsic HR (iHR) (11, 12). Notably, circadian rhythms in iHR can be eliminated with I_HCN_ blockade, confirming the essential role of I_HCN_ in mediating LCR within the sinoatrial node (8).

Regulated jointly by the ANS and LCR (Fig 1), circadian rhythms in HR display intriguing properties of both flexibility and robustness. They exhibit flexibility, allowing for a wide range of pacing frequencies, and optimizing time-of-day cardiac outputs under diverse physiological conditions, such as sleep/awake or inactive/active states. Simultaneously, these rhythms remain robust, ensuring the generation of rhythmic HR patterns throughout the day despite changes in external conditions. However, how the ANS and LCR interact to facilitate circadian flexibility in cardiac pacemaking function with robustness remains unclear. Furthermore, their vulnerability to severe perturbations, such as aging, remains enigmatic (Fig 1; red dot) (13-15).

To unravel these questions, the application of mathematical modeling and simulation proves invaluable in providing mechanistic insights into the non-linear behaviors of complex biological systems, e.g., mammalian circadian dynamics (16-20) and cardiac excitation patterns (21, 22). Earlier *in silico* studies have quantified the impact of circadian expression of potassium channel interacting protein-2 (KChIP2) on shaping ventricular action potential (AP) morphologies (23, 24), and investigated the role of circadian rhythmicity of I_CaL_ expression and function in the occurrence of early after-depolarizations (EAD) in guinea pig ventricular cardiomyocytes and tissues (25). However, previous computational studies of sinoatrial nodal cells (SANC) have mainly emphasized the ionic interactions between Ca^2+^ and membrane oscillations in the generation of spontaneous pacemaking activities (22, 26). There remains a paucity of *in silico* studies that aim to comprehend the circadian regulation of cardiac pacemaking function. In particular, the central and local circadian aspects of SANC automaticity have not been established.In this study, we present a novel mathematical model that captures the intricate regulation of SANC by both autonomic control and LCR in mice (Fig 1). The model accurately reproduces diverse circadian patterns as shown in previous experimental studies in adult and aged mice (7, 8, 14).

Utilizing the model, we elucidate the specific roles of PNA, SNA, and LCR in optimizing circadian flexibility and robustness in SANC, as well as quantitatively dissecting SANC dysfunction during the aging process. Specifically, SNA serves as a SANC robustness and performance enhancer, PNA acts as a flexibility amplifier, and LCR functions as a flexibility and performance booster. However, with aging, SANC flexibility is dampened, and mainly influenced by LCR attributable to aging-related ion channel remodeling and PNA impairment. Our model simulations highlight a critical dimension for time-of-day interactions between PNA, SNA, and LCR in cardiac pacemaking function and dysfunction.

## Results

### Quantitative reconstruction of diverse circadian patterns in SANC FR

It has been suggested that in anesthetized mice, PNA is more dominant in determining circadian HR variations compared to SNA (Fig 2A; blue vs yellow dots) (7). Specifically, the amplitude of circadian heart rate (HR) variations showed a significant reduction of 75% with a PNA blockade (PNAB), whereas an SNA blockade (SNAB) resulted in a smaller reduction of 16%. Moreover, intriguingly, even after a complete autonomic nervous system blockade (ANSB), circadian HR variations persisted (Fig 2A; green dots). This suggests that there are other factors or mechanisms involved in regulating the circadian rhythmicity of heart rate in anesthetized mice, in addition to the ANS activity.

**Fig 2.**
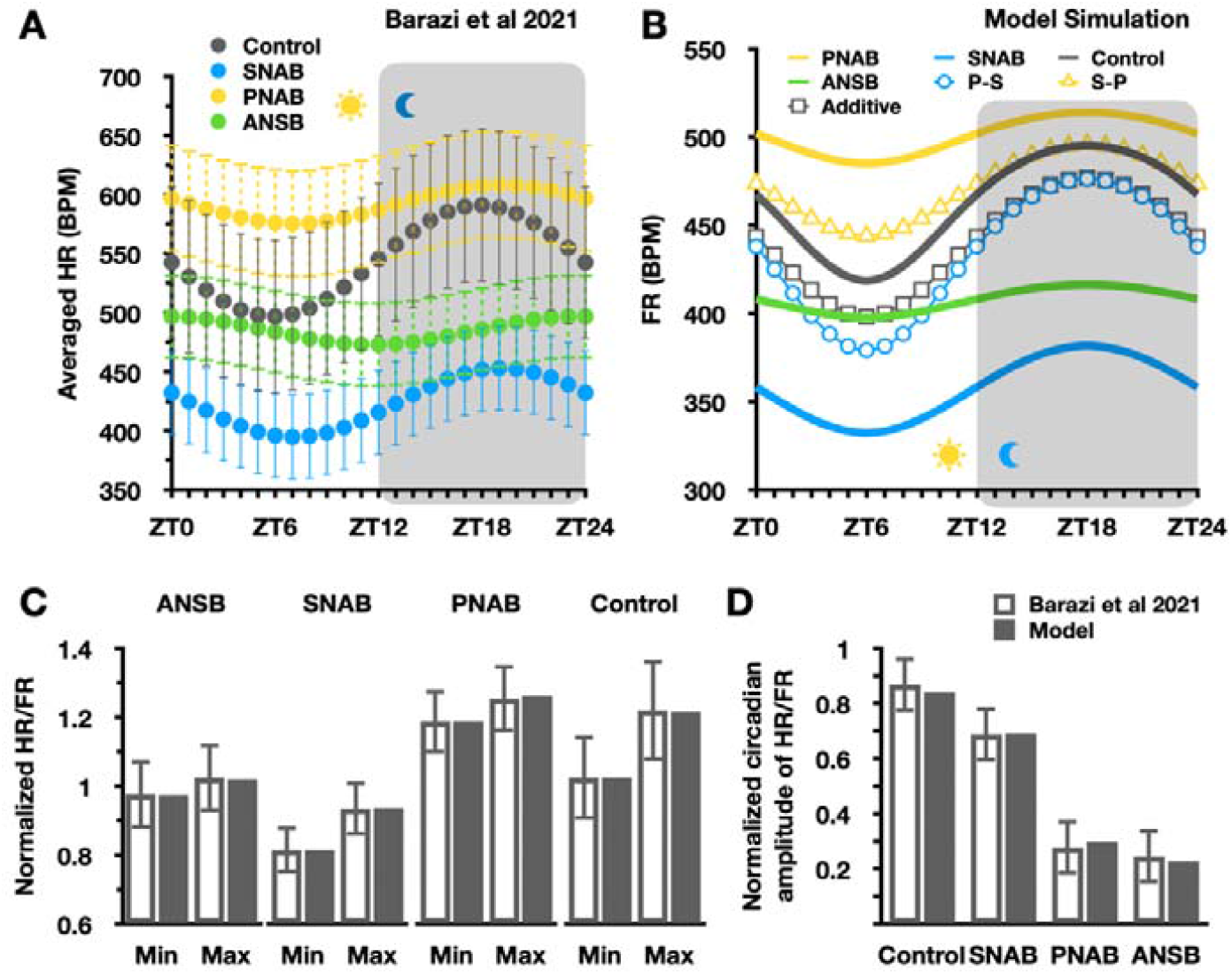
Quantitative reconstruction of diverse circadian patterns in SANC FR under various conditions. (A) Circadian HR fluctuations in anesthetized mice with 12-h light cycles before (control; grey dots) and after SNAB with propranolol (blue dots), PNAB with atropine (yellow dots), and ANSB with atropine + propranolol (green dots) (7). (B) Simulated circadian patterns in mouse SANC FR closely recapitulate experimental findings (7). Specifically, LCR alone generates a minor circadian amplitude of FR with ANSB (green line), similar to what is observed in (A; green dots). The circadian amplitude of FR is marginally increased with PNAB (LCR+SNA) (yellow line), yet it is significantly enhanced with SNAB (LCR+PNA) (blue line), as experimentally observed in (A; yellow and blue dots). However, when assuming additive PNA and SNA (no PNA-SNA interactions; grey box), simulated FR deviates from the experimental data under the control conditions. On the other hand, the incorporation of bidirectional modulation effects (non-additive PNA and SNA; grey line) aligns the simulated FR with experimental data (A; grey dots). Furthermore, unidirectional modulation effects alone (S-P (yellow triangle) or P-S (blue circle)) are both insufficient to accurately reproduce circadian patterns under the control conditions. (C) Model simulations accurately reproduce both minimal and maximal time-of-day FR under control, SNAB, PNAB, and ANSB conditions. The FR values are normalized to FR at ZT12 under the ANSB condition, providing a quantitative comparison across the different experimental setups. (D) Model simulations precisely replicate normalized circadian amplitudes of FR under control, SNAB, PNAB, and ANSB conditions. The normalized amplitude values are obtained by dividing circadian FR amplitudes in BPM by the mean time-of-day FR.

Our model simulations (Fig 2B) meticulously recapitulated these key experimental findings under control, SNAB, PNAB, and ANSB conditions with normalized circadian amplitudes of FR (Fig 2D). Specifically, when assuming additive PNA and SNA (no PNA-SNA interactions), the simulated circadian rhythm of normalized FR exhibits a similar amplitude compared to the experimental data. However, the baseline of the simulated circadian rhythm of normalized FR (Fig 2B; grey box) is lower by ∼5% in comparison to the experimental data (Fig 2A; grey dots). In contrast, when we implemented bidirectional (PNA to SNA, P-S; SNA to PNA, S-P) modulation effects in our model to adopt their non-additive interactions (see Method and Supporting Information for details), the circadian pattern in FR under the control condition was accurately simulated (Fig 2B; grey line) (Fig 2C) (6, 7). Specifically, the P-S modulation alone tends to enhance the circadian amplitude of FR (Fig 2B; blue circle), while the S-P modulation alone elevates its baseline (Fig 2B; yellow triangle). As a result of combined P-S and S-P effects, both minimal and maximal time-of-day FR are in quantitative agreement with the experimental data (Fig 2C). Moreover, the simulated circadian amplitude of FR is also well aligned with the experiments (Fig 2D). Our results unequivocally indicate that the non-additivity of PNA and SNA is required to properly reconstruct the circadian patterns of SANC function.

Despite the circadian amplitude of FR being largely preserved with SNAB compared to the control condition (Fig 2B), the presence of non-additive interactions between PNA and SNA leads to distinct mechanisms by which PNA regulates the circadian amplitude of FR. These mechanisms are contingent on the presence or absence of SNA. In the absence of SNA, PNA directly regulates the circadian amplitude in FR through its effect on I_KACh_ alone (Fig 1). Conversely, in the presence of SNA, PNA regulates circadian amplitude in FR via indirect time-of-day “braking” effects on the SNA as well as its direct regulation of I_KACh_ (see Supporting Information for details).

### The synergistic combination of SNA, PNA, and LCR is essential in optimizing the time-of-day SANC function

Our model successfully recapitulated diverse circadian patterns in SANC pacemaking function as shown in Fig 2. Subsequently, we utilized the model to investigate how SNA, PNA, and LCR interact to maintain circadian flexibility in SANC pacemaking function with robustness and performance. To achieve this, we simulated Ca^2+^ oscillator (CO) – membrane oscillator (MO) parameter space maps at ZT6 and ZT18 under control, SNSB, PNSB, ANSB, and LCR blockade (LCB) conditions (Fig 3). Specifically, each CO-MO parameter space map was generated by reducing the maximal rate of SERCA (P_up_) (a CO parameter) and the maximal conductance of I_CaL_ and I_CaT_ (MO parameters) from their original values to zero. The original values for these CO and MO parameters were defined as previously described (27), with P_up_= 0.04 mM/ms, G_CaL_= 0.018 nS/pF and G_CaT_ = 0.013956 nS/pF. The steady-state FR in BPM was computed for each simulation and color-coded to create the map. In the CO-MO parameter space maps, we quantified SANC robustness as the percentage area of rhythmic firing regions, which indicates the regions in the parameter space where SANC maintains rhythmic pacemaking. Borders between rhythmic firing regions and chaotic or no firing regions were identified by detecting the nearest local trough from the original values of CO and MO (Fig 3A; Control, blue lines). Under control conditions, SANC robustness is 58.7% and 54.3% at ZT6 and ZT18, respectively (Fig 3A; Control) (Fig 3B). SANC flexibility was measured as the difference between minimal (ZT6) and maximal (ZT18) time-of-day SANC FR with control CO and MO parameter values, reflecting the ability of SANC to adjust its FR over the circadian cycle. SANC flexibility under control conditions is 76.7 BPM (Fig 3C), which is the difference between 495.3 BPM (ZT18) and 418.6 BPM (ZT6) (Fig 3A). In addition, SANC performance was assessed as the maximal time-of-day SANC FR, representing the peak FR reached during the circadian cycle, which is 495.3 BPM in the control (Fig 3D).

**Fig 3.**
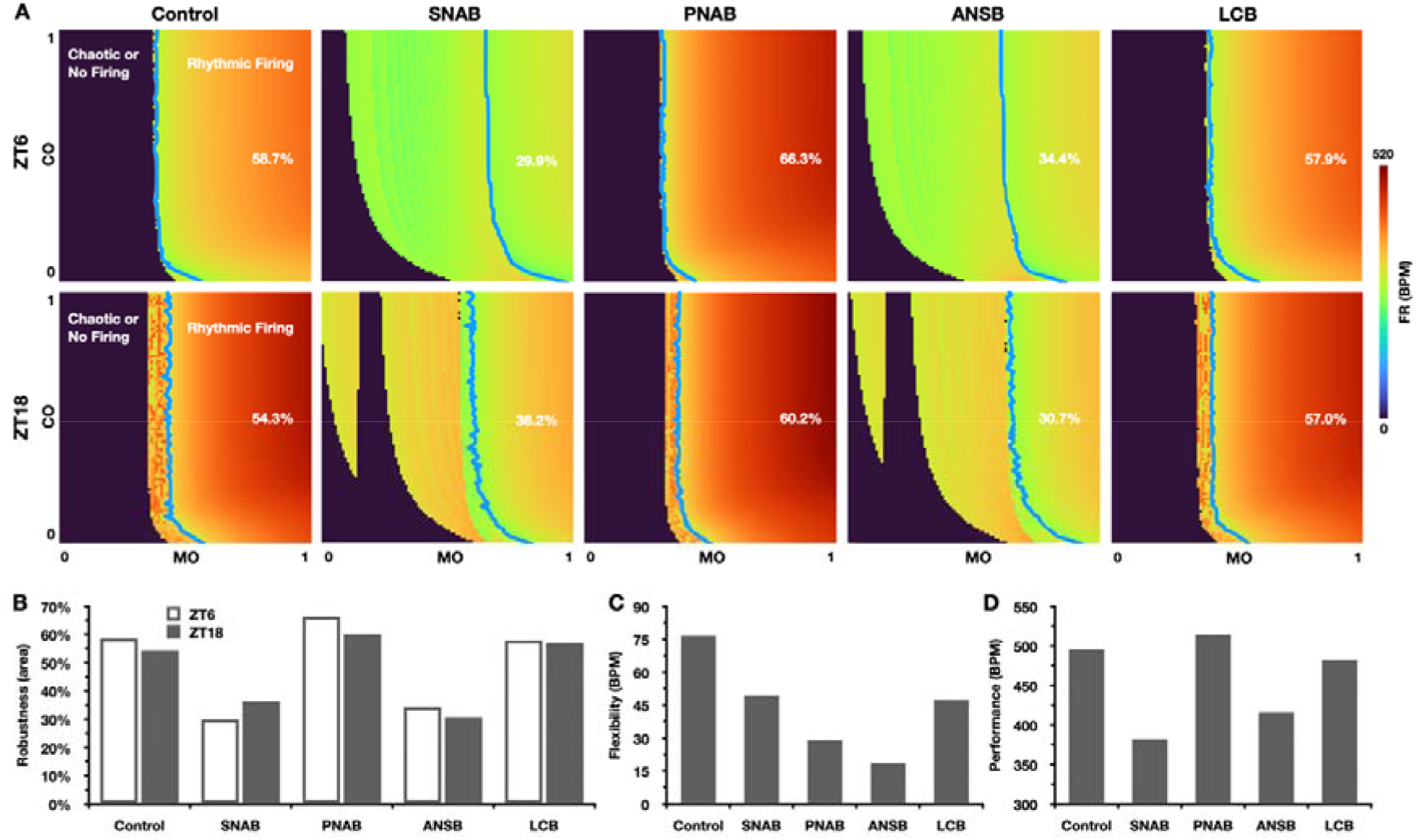
The synergistic combination of SNA, PNA, and LCR is essential to achieve circadian flexibility with robustness and performance in SANC automaticity. (A) CO-MO parameter space maps color-coded by FR in BPM at ZT6 and ZT18 under control, SNAB, PNAB, ANSB, and LCB conditions. In each CO-MO parameter space map, the maximal rate of SERCA (P_up_) (CO; along the vertical axis), and the maximal conductance of I_CaL_ and I_CaT_ (MO; along the horizontal axis) were reduced from their original values to zero with a step size of 1%, resulting in a total of 100 × 100 simulations. Each simulation was color-coded by FR in BPM. Blue solid lines demarcate the border between rhythmic firing and chaotic or no-firing regions. (B-D) Quantitative differences in robustness (B), flexibility (C), and performance (D) under various conditions. While SNA enhances SANC robustness and performance, PNA magnifies SANC flexibility. LCR acts as a booster for SANC flexibility and performance. With the synergistic combination of SNA, PNA, and LCR, SANC exhibits maximal circadian flexibility (76.7 BPM) with both robustness (58.7% and 54.3% at ZT6 and ZT18 respectively) and performance (495.3 BPM).

When SNA is inhibited, SANC robustness (29.9% and 36.2% at ZT6 and ZT18, respectively) (Fig 3A; SNAB) (Fig 3B) and performance (381.9 BPM) (Fig 3D) are significantly reduced, with a moderate impact on SANC flexibility (49.5 BPM) (Fig 3C). In contrast, the blockade of PNA enhances both SANC robustness (66.3% and 60.2% at ZT6 and ZT18, respectively) (Fig 3A; PNAB) (Fig 3B) and performance (514.1 BPM) (Fig 3D), but there is a notable reduction in SANC flexibility (29.1 BPM) (Fig 3C). Under ANSB, all aspects of SANC pacemaking function are impaired, with residual circadian flexibility of 18.6 BPM. Meanwhile, the LCR blockade has a relatively minor impact on SANC robustness (57.9% and 57.0% at ZT6 and ZT18, respectively) compared to the control (Fig 3A; ANSB) (Fig 3B). However, SANC flexibility is markedly reduced to 47.4 BPM (Fig 3C), along with a moderate reduction in SANC performance to 482.3 BPM (Fig 3D). These findings indicate that SNA, PNA, and LCR each play distinct roles in shaping circadian patterns in SANC FR: SNA acts as a robustness and performance enhancer, PNA acts as a flexibility magnifier, and LCR acts as a booster for both flexibility and performance. Furthermore, these factors synergistically interact to achieve high circadian flexibility with robustness and performance in SANC pacemaking function (Fig 3B-3D).

### Quantitative dissection of SANC pacemaking dysfunction in aging

Age-related PNA impairment has been documented since the early stage of aging in mice (15, 28). In addition, aging is also characterized by a progressive decline in maximal HR, attributed to aging-dependent MO reduction (i.e., ion channel remodeling in I_CaT_, I_CaL_, and I_HCN_) (14). We developed an aging model that can successfully simulate the experimentally measured aging-dependent PNA impairment and MO reduction (14, 15, 28) (see Method and Supporting Information for details). Then, to understand how the ANS and LCR fail to maintain SANC pacemaking function under aging, we conducted simulations of CO-MO parameter space maps at ZT6 and ZT18 under control, aging-dependent PNA impairment or/and MO reduction (Fig 4). When considering only PNA impairment, SANC robustness remained mostly unchanged (60.2% and 52.9% at ZT6 and ZT18, respectively) (Fig 4A; ↓PNA) (Fig 4B), while SANC flexibility was dampened from 76.7 BPM to 37.6 BPM (Fig 4C), with a minor decrease in SANC performance from 495.3 BPM to 483.4 BPM (Fig 4D). In contrast, with aging-dependent MO reduction alone, SANC robustness was preferentially reduced at ZT6 compared to ZT18 (Fig 4A; ↓MO) (Fig 4B), decreasing from 58.7% to 46.8%, and SANC flexibility and performance were reduced to 50.5 BPM and 356.8 BPM, respectively (Fig 4C-4D). When aging conditions were combined, the decline in SANC flexibility and performance escalated to 16.9 BPM and 347 BPM (Fig 4C-4D), respectively. Notably, SANC robustness at ZT6 (the sleep/inactive phase) improved from 46.8% to 56.3% compared to the sole MO reduction condition (Fig 4A; Aging) (Fig 4B). Our findings indicate a potential time-of-day difference in SANC dysfunction associated with aging. Specifically, the inactive/sleep phase (e.g., at ZT6) might exhibit greater impairment due to aging-related ion channel remodeling compared to the active/awake phase (e.g., at ZT18).

**Fig 4.**
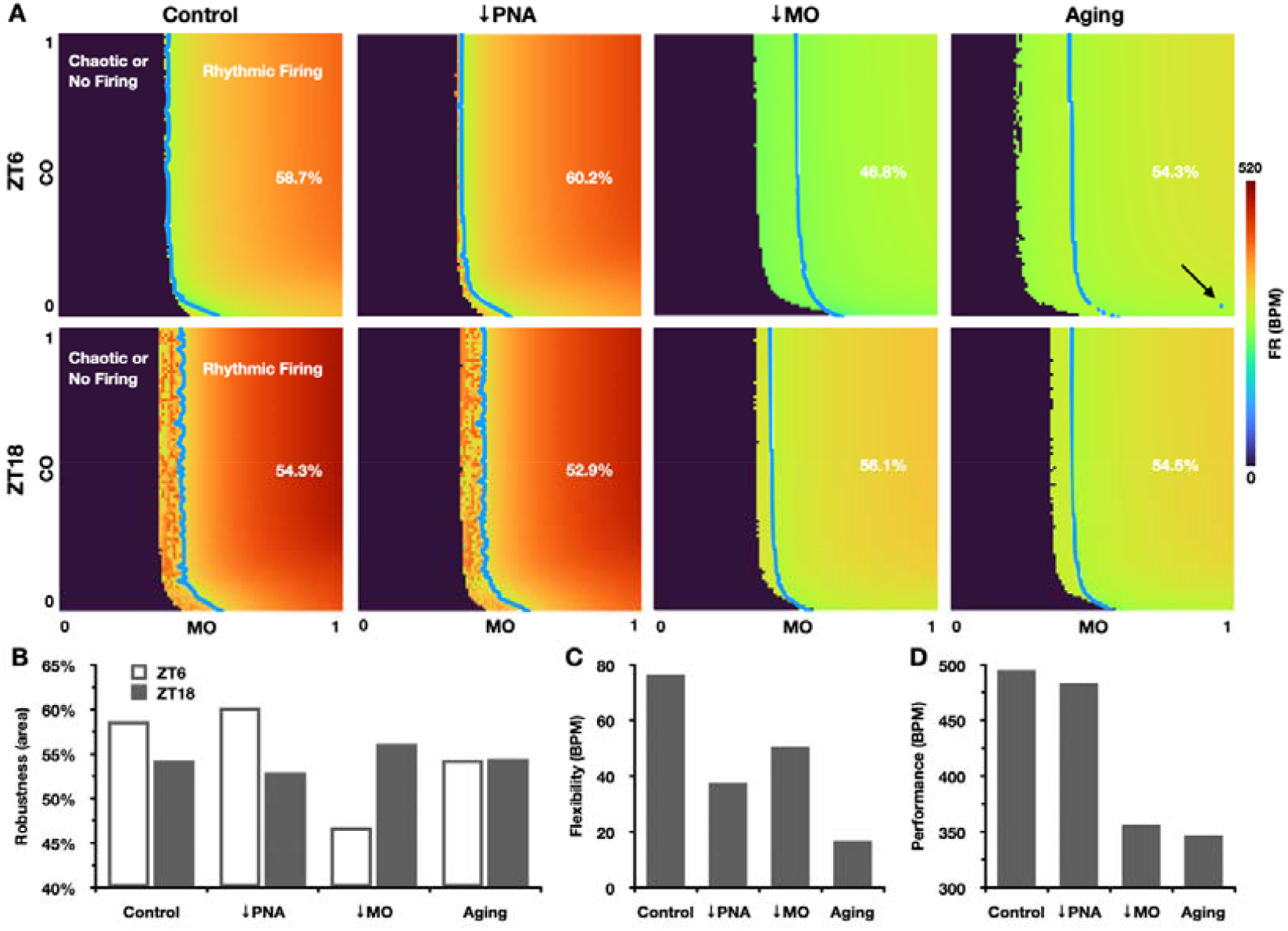
Quantitative dissection of SANC pacemaking dysfunction in aging. (A) CO-MO parameter space maps at ZT6 and ZT18 under control, PNA impairment, MO reduction, and full aging conditions. Blue solid lines define the border between rhythmic firing and chaotic or no-firing regions. The arrow indicates the incidence of SANC pacemaking instability. (B-D) Quantitative differences in robustness (B), flexibility (C), and performance (D) under various aging conditions. While MO reduction has a detrimental effect on all aspects of SANC function, PNA impairment preferentially reduces SANC flexibility with a protective role in SANC robustness at ZT6. Full aging effects are characterized by minimal SANC flexibility (16.7 BPM) and performance (347 BPM) with preserved robustness.

### Distinct mechanisms underlying circadian patterns in SANC FR in adult and aged mice

To dissect key mechanisms underlying circadian patterns in SANC, we conducted a numerical analysis to assess and compare the relative contributions of each module in the model in determining circadian patterns of SANC. This assessment was quantified by the extent to which either flexibility or performance of SANC was perturbed by fully inhibiting PNA, SNA, central circadian rhythmicity (CCR), LCR, P-S or S-P modulation effects (Fig 5A; gray). Here, CCR refers to the circadian component of ANS, and specifically, CCR is the circadian component of PNA given that SNA is modeled with no circadian variations in our study. In adult mice (Fig 5A; grey), while PNA (62%) dominates the circadian flexibility of SANC, LCR (38%) and SNA (35%) play a secondary role in promoting circadian flexibility. The role of SNA in SANC flexibility is counterintuitive given that SNA is modeled with no circadian fluctuations. Such observation is attributable to the PNA-dependent “braking” effects of SNA (P-S; Fig 2B, blue circle). Moreover, we found that unidirectional P-S (32%) and S-P (−27%) modulation effects alone exhibit opposite effects on the circadian flexibility of SANC (Fig 5A; grey). The role of [Na]_i_^+^ homeostasis in regulating cardiac excitation patterns has been previously reported (29-31). To evaluate its potential role in regulating circadian patterns in SANC, we clamped the time-of-day [Na]_i_^+^ content at its steady state concentration at ZT6. Our findings revealed that [Na]_i_^+^ homeostasis exerts a dampening effect (−10%), which can be attributable to a higher [Na]_i_^+^ content at faster FR (Fig 5A; grey). As expected, SNA and PNA are most important in enhancing or decreasing SANC performance, respectively. In Fig 5B (grey), we conducted a sensitivity analysis to quantify the individual contributions of key model parameters in shaping circadian patterns of SANC. Each parameter value was reduced by 25% to determine its impact. In adult mice, I_CaT_ and I_Kr_ are the most important contributors in promoting or dampening SANC flexibility and performance, respectively.

**Fig 5.**
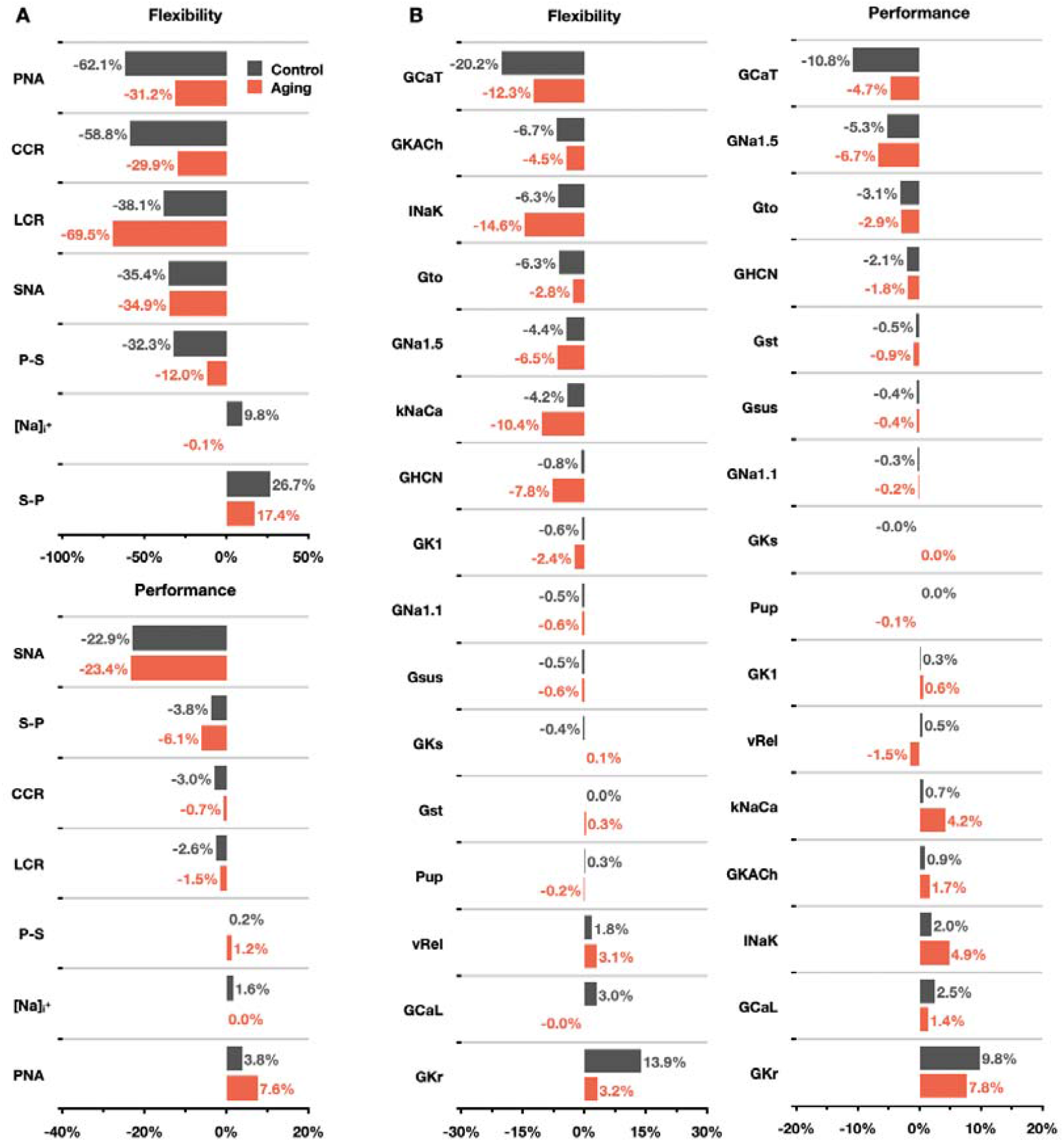
Distinct mechanisms underlying circadian patterns of SANC FR in control and aged mice. (A) Relative contributions to SANC flexibility and performance are quantified by selective inhibition of each module in control (grey) and aging (red) conditions. The relative contributions of PNA, SNA, P-S and S-P modulation effects were assessed through complete inhibition of their activities. To inhibit CCR and LCR, the circadian amplitudes of ANS and I_HCN_ were reduced to zero, respectively. Additionally, the influence of [Na]_i_^+^ homeostasis was evaluated by clamping time-of-day [Na]_i_^+^ content at its steady state concentration at ZT6. (B) Sensitivity analysis of SANC flexibility and performance was conducted by applying a 25% reduction to key model parameter values in control (grey) and aging (red) conditions. Full model parameter definitions are provided in the Supporting Information S1 Table.

However, in aged mice, distinct mechanisms underlying the circadian flexibility of SANC were observed in Fig 5A (red). The circadian amplitude of SANC in aging is mainly influenced by LCR (70%), attributable to aging-related ion channel remodeling and PNA impairment. Additionally, SNA (35%) and PNA (31%) make secondary contributions to circadian amplitude, while [Na]_i_^+^ homeostasis plays a negligible role. In Fig 5B (red), due to aging-related ion channel remodeling, the role of I_NaK_, I_NaCa_ becomes more substantial in preserving SANC flexibility and reducing SANC performance.

## Discussion

In this study, we developed a model of autonomic control and LCR in SANC to recapitulate diverse circadian patterns in both adult and aged mice. Utilizing this model, we elucidated the distinct roles of PNA, SNA, and LCR in optimizing circadian flexibility with robustness and performance in SANC, as a flexibility amplifier, an enhancer for robustness and performance, and a booster for flexibility and performance, respectively. Specifically, while SANC robustness and performance are preferentially enhanced by SNA, SANC flexibility is primarily driven by PNA with a secondary contribution of LCR and SNA, and moderately dampened by [Na]_i_^+^ homeostasis. The secondary role of SNA in SANC flexibility could be counterintuitive given that SNA is modeled with no circadian variations in our study, and it is attributable to the PNA-dependent “braking” effects of SNA. Compared to PNA and SNA, LCR acts as a booster for SANC flexibility and performance, with limited effects on robustness. Due to the distinct roles of PNA, SNA, and LCR, they synergistically work together to achieve circadian flexibility with robustness and performance in SANC. However, this synergy is broken under aging conditions, leading to a reduction in the performance and flexibility of SANC. Specifically, aging-dependent PNA impairment selectively dampens SANC flexibility with a protective role in preserving SANC robustness. As a result of aging-related ion channel remodeling (i.e., reduction in I_CaT_, I_CaL_, I_HCN_), all aspects of SANC function (i.e., robustness, flexibility, and performance) are disrupted.

It is known that circadian rhythms in HR can arise from either the master circadian clock located in the SCN or the local circadian clock in the heart (2, 4). Previous experimental studies have suggested that the 24h HR rhythm is primarily governed by the SCN through the ANS (4, 8, 32, 33). However, instead of solely focusing on determining which one, the master or local circadian clock, dominates, our study was motivated to understand the necessity of having two circadian clocks to regulate cardiac pacemaking function and to explore their specific roles. Earlier computational studies have primarily focused on the dynamic interactions between Ca^2+^ and membrane oscillations in generating spontaneous pacemaking activities in SANC (26, 34). However, the circadian aspects of SANC pacemaking function over the course of the day have not been established (35). Our study sheds light on a critical yet understudied dimension of time-of-day interactions between PNA, SNA, and LCR in cardiac pacemaking function and dysfunction. As illustrated in Fig 6A, after departing up-hill from the baseline state (grey circle), LCR (green circle) introduces moderate SANC flexibility and a slight increase in performance without affecting robustness. The addition of SNA (yellow circle) primarily enhances SANC robustness and performance, while PNA (blue circle) promotes SANC flexibility at the expense of some performance. The synergistic combination of SNA, PNA, and LCR ultimately leads to the baseline-to-control up-hill trajectory, achieving high levels of robustness, flexibility, and performance (grey dot). LCR serves as an important booster for SANC flexibility and performance (green dot), and its extra, boosting effects may offer a survival advantage when confronted with external threats. In Fig 6B, after departing down-hill from the control state, the reduction of MO in aging undermines all aspects of the SANC function, while aging-dependent PNA impairment preferentially reduces SANC flexibility with little effect on robustness and performance. With combined aging effects, both SANC performance and flexibility were more impaired than with any aging effect alone. However, SANC robustness remained relatively preserved, indicating a potential “trade-off” strategy where flexibility and performance are sacrificed to ensure safety. This observation aligns with previous experimental findings indicating age-related attenuation of parasympathetic control of the heart in mice (28).

**Fig 6.**
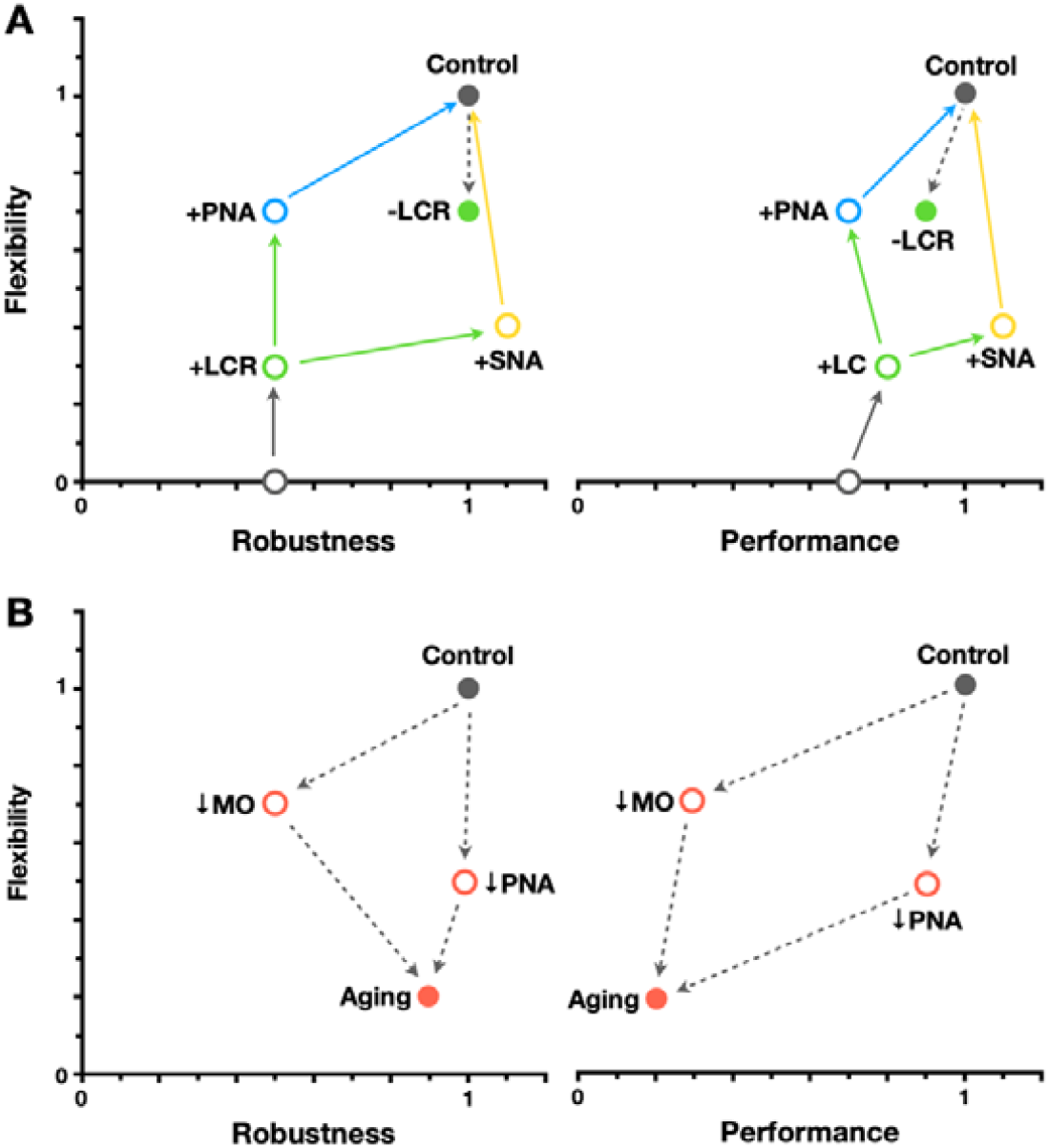
Illustrative trajectories in the parameter space of SANC robustness, flexibility, and performance. (A) Up-hill trajectories show the transition from baseline (ANSB + LCB; grey circle) to control (ANS + LCR; grey dot) model states. (B) Down-hill trajectories demonstrate the transition from control (grey dot) to aging model states (pink circles and dots). SANC robustness is defined as the normalized minimal time-of-day area percentage of the rhythmic firing region, flexibility as the normalized difference between SANC FR at ZT6 and ZT18, and performance as the maximal time-of-day SANC FR.

Our control model was calibrated to experimental data in anesthetized mice (7) assuming limited circadian activities in SNA (15), which may potentially underestimate the role of SNA in SANC flexibility in free-moving mice living in a thermoneutral environment (36). Our aging model was parameterized based on a cross-sectional experimental dataset in >32 months old mice, thus lacking the capacity to capture functional trajectories throughout the aging process. We modeled LCR as a diurnal variation in the expression level of I_HCN_, but it might not be the only mechanism underlying LCR in SANC. For example, a day/night difference has been reported in Ca^2+^/calmodulin-dependent protein kinase II delta (CaMKIIδ) expression (7). In addition, our model is limited by the lack of a biochemical description of beta-adrenergic and cholinergic signaling pathways and their interactions, and a detailed TTFL model for LCR in SANC, due to the scope of this study and the limited availability of experimental data. Additional model development would be helpful to further advance our understanding of circadian SANC function and dysfunction, e.g., the mechanistic coupling between the ANS and LCR (37), trajectory of functional aging (38), and entrainment dynamics between the master circadian clock and LCR during jet lag or with shift work disorder (39).

## Materials and Methods

### Model development

A mathematical model of mouse SANC AP (27, 31, 40) was used as our baseline model (without the ANS and LCR) (Fig 1). LCR was then introduced into the model as a circadian variation in the expression and function level of I_HCN_ (Fig 1; green dot). This addition resulted in a minor (∼2%) circadian amplitude in SANC firing rates (FR) as observed in immobilized anesthetized mice under full ANS blockade conditions, with its trough occurring at zeitgeber time (ZT) 6 (7, 8) (Fig 2A; green dots). A 12h:12h light/dark lighting regime was implemented as in the experimental studies (7, 8). Time-of-day PNA (Fig 1; blue dot) was then incorporated and calibrated as a circadian function of carbachol (CCh) concentrations using model formulations previously described (27), based on experimental measurements of normalized mean HR and circadian amplitude of HR under SNAB conditions (7) (Fig 2A; blue dots). Similarly, time-of-day SNA (Fig 1; yellow dot) was implemented and calibrated using earlier approaches to model isoproterenol (ISO) effects (27), based on experimental data under PNAB conditions (7) (Fig 2A; yellow dots). We noted that a combination of LCR and ISO effects (27) would be enough to generate a circadian amplitude (∼3%) in SANC FR as measured in the experiments (7) and leave little room for additional in-phase circadian variations in SNA/ISO. Consequently, diurnal SNA was modeled as a high sympathetic tone with little circadian variation. This modeling choice aligns with earlier experimental findings on the effects of a thermoneutral environment on heart rate variability, where mice exhibit a high sympathetic drive to maintain a normal core temperature (37°C) under standard laboratory conditions (20°C) (15, 36).

Under control conditions, bidirectional modulation effects between PNA and SNA (P-S and S-P) (Fig 2B) were formulated to account for the non-additivity effects as previously reported (6, 41-43), and calibrated to be consistent with experimental measurements (7). Aging-dependent MO reduction (Fig 1; red dot), i.e., ion channel remodeling in I_CaT_, I_CaL_, and I_HCN_, was implemented and calibrated to be consistent with electrophysiological data in aged mice (>32 months old) (14). Aging-dependent PNA impairment (28) (Fig 1; red dot) was introduced by reducing the circadian amplitude of PNA and calibrated to reproduce a major reduction in the maximal HR (at ZT18) as experimentally observed in aged mice (14). Full model details are provided in the Supporting Information II and S2 Table.

### Simulation protocols

Circadian patterns of ANS and LCR were considered to be in phase with each other and aligned at ZT6 (Fig 2B). Steady-state FR was calculated as the averaged FR of the last 10 spontaneous AP after 60 seconds of simulation. Each CO-MO parameter space map (26) was reproduced by reducing the maximal rate of SERCA (P_up_) (CO) and the maximal conductance of I_CaL_ and I_CaT_ (MO) from their original values (27) to zero, using a step size of 1%, resulting in a total of 100 × 100 simulations. Each simulation was color-coded by steady-state FR in BPM. Borders between rhythmic firing regions and chaotic or no firing regions were identified by detecting the nearest local trough from the original values of CO and MO. To quantify SANC robustness using CO-MO parameter space maps (Fig 3-4), we define robustness as the percentage area of rhythmic firing regions. To quantify SANC flexibility and performance, we define flexibility as the difference between the minimal and maximal time-of-day SANC FR in BPM with the control CO and MO, and performance as the maximal time-of-day SANC FR in BPM. Sensitivity analysis was conducted with either a full inhibition or a reduction (25%) of each module or parameter value of interest (Fig 5). Additionally, sensitivity analysis for [Na]_i_^+^ homeostasis was performed by clamping time-of-day [Na]_i_^+^ content at its steady state concentration at ZT6.

## Acknowledgments

The authors thank all members of the Biomedical Mathematics Group at the Institute for Basic Science for their most helpful discussions.

